# Functional and Structural Moment Arm Validation for Musculoskeletal Models: A Study of the Human Forearm and Hand

**DOI:** 10.1101/2020.05.29.124644

**Authors:** Matthew T Boots, Russell Hardesty, Anton Sobinov, Valeriya Gritsenko, Jennifer L. Collinger, Lee E. Fisher, Robert Gaunt, Sergiy Yakovenko

## Abstract

The development of realistic musculoskeletal models is a fundamental goal for the theoretical progress in sensorimotor control and its engineering applications, e.g., in the biomimetic control of artificial limbs. Yet, accurate models require extensive experimental measures to validate structural and functional parameters describing muscle state over the full physiological range of motion. However, available experimental measurements of, for example, muscle moment arms are sparse and often disparate. Validation of these models is not trivial because of the highly complex anatomy and behavior of human limbs. In this study, we developed a method to validate and scale kinematic muscle parameters using posture-dependent moment arm profiles, isometric force measurements, and a computational detection of assembly errors. We used a previously published model with 18 degrees of freedom (DOFs) and 32 musculotendon actuators with force generated from a Hill-type muscle model. The muscle path from origin to insertion with wrapping geometry was used to model the muscle lengths and moment arms. We simulated moment arm profiles across the full physiological range of motion and compared them to an assembled dataset of published and merged experimental profiles. The muscle paths were adjusted using custom metrics based on root-mean-square and correlation coefficient of the difference between simulated and experimental values. Since the available measurements were sparse and the examination of high-dimensional muscles is challenging, we developed analyses to identify common failures, i.e., moment arm functional flipping due to the sign reversal in simulated moments and the imbalance of force generation between antagonistic groups in postural extremes. The validated model was used to evaluate the expected errors in torque generation with the assumption of constant instead of the posture-dependent moment arms at the wrist flexion-extension DOF, which is the critical joint in our model with the largest number of crossing muscles. We found that there was a reduction of joint torques by about 35% in the extreme quartiles of the wrist DOF. The use of realistic musculoskeletal models is essential for the reconstruction of hand dynamics. These models may improve the understanding of muscle actions and help in the design and control of artificial limbs in future applications.

**New & Noteworthy:** Realistic models of human limbs are a development goal required for the understanding of motor control and its applications in biomedical fields. However, developing accurate models is restrained by the lack of accurate and reliable musculoskeletal measurements in humans. Here, we have overcome this challenge by using multi-stage validation of simulated structures using both experimental data and the identification of structural failures in the high-dimensional muscle paths. We demonstrate that the rigorous structural and functional validation method is essential for the understanding of force generation at the wrist.

## Introduction

Movement is a fundamental behavior of all living organisms allowing them to manipulate the external environment to achieve survival objectives. The motor system is easily able to coordinate complex muscle activation patterns to produce torques that lead to skilled movements. Extreme examples are human thumb muscles representing high-dimensional kinematic relationships as a function of six degrees of freedom (DOF). It is not surprising that thumb control is a challenge in powered prosthetics {ref} or in the related process of decoding motor intent from electromyography {ref}. The sophisticated algorithms for the control of relatively simpler artificial limbs have yet to reach the robustness and accuracy of their biological counterparts (Crouch and Huang, 2016; Dantas et al., 2019; Resnik et al., 2018). One biomimetic solution is the model-based prosthetic control mimicking the presence of MS computations within neural computations of planning and execution pathways (Lillicrap and Scott, 2013; Shadmehr et al., 2016). In engineering, the use of embedded models of controlled structure has been employed in the Smith’s predictor approach (Smith, 1957) and more recently, non-linear system models have been used in the model predictive control applications. This general approach can be potentially used for prosthetic control (Crouch and Huang, 2016; Sartori et al., 2018); however, achieving robust control with the multidimensional MS models is the challenge, especially in real-time applications.

The effort to create valid muscle-driven MS models spans dozens of years in the context of non-invasive analysis of gait, posture, and reaching movements (Arnold et al., 2010; Carbone et al., 2015; De Pieri et al., 2018; Delp et al., 1990; Gritsenko et al., 2016; Horsman, 2007; Rajagopal et al., 2016; Saul et al., 2015b). The models are typically developed using minimalistic sets of parameters and with additional testing of model performance against experimental observations (Kirchner, 2006). The popular truism, “*all models are wrong, but some are useful,*” expressed by statistician George Box (Box, 1979), guides the best practice of developing models and their validation in ever more ambitious biomechanical analyses. However, the main limiting factor in model development is the sparseness of experimental structural and functional datasets. We have addressed here three common limitations of MS models that limit their robustness: 1) errors in simulated muscle path for experimentally observed postures; 2) errors in simulated muscle path for experimentally unobserved postures; 3) force scaling mismatch between muscle groups.

In general, MS models are tested through either direct or indirect validation (Henninger et al., 2010; Lund et al., 2012). Comparing simulated and measured muscle attributes like moment arms is an example of a direct morphological validation (Arnold et al., 2001, 2000; Delp et al., 1990; Holzbaur et al., 2005). In the indirect validation methods, the musculoskeletal attributes are initially assigned based on the experimental or computational data, but their validation is achieved through the demonstration of matching profiles in simulated and recorded muscle activity patterns or their forces (de Zee et al., 2007; Hamner et al., 2010). These signals are computed from the sequence of computations starting with inverse kinematics to compute joint angles. These are then passed through inverse dynamics to compute joint torques. The last step is the constrained computation for the ill-posed problem due to the muscle redundancy where the number of joint torques is exceeded by the number of muscle moments that contribute to these torques (Crowninshield and Brand, 1981; Erdemir et al., 2007). A direct comparison is preferred; however, there is limited availability of moment arm data for different postures. Often, MS models rely on disparate data sources, i.e., measurements combined across cadavers and different studies that may use different methodologies. The combined models may inherit inter-subject variations based on measurements that are not correctly scaled and lead to unphysiological nonlinearities (Goislard De Monsabert et al., 2018). This problem necessitates the use of additional indirect validations that examine overall function. For example, scaling the forces of individual muscles by either changing their force generation parameters (Scovil and Ronsky, 2006) or their muscle moment arms (Nussbaum et al., 1995) to match observed torque measurements.

In this study, we aimed to overcome structural and functional model inaccuracies by applying a novel direct structural validation method combined with an indirect validation of functional output. The main focus was to develop a validation method that can overcome the previous limitations in this model (Gritsenko et al., 2016; Saul et al., 2015a) with the sparseness and disparity of experimental measurements. We then quantified the contribution of nonlinear moment arm profiles to the generation of joint moments compared to those produced with constant moment arms to test the necessity for dynamic moment arm.

## Methods

### Overview

The model validation process consisted of the following three steps: *i*) the creation of a meta dataset describing muscle moment arm values for a sparse selection of postures from published studies, *ii*) the selection of wrapping geometry to constrain muscle paths, and *iii*) the validation of structural and functional muscle properties.

### Model

We used a model of an arm and hand developed by Saul et al. specifically as a platform for future development and benchmarking (Gritsenko et al., 2016; Saul et al., 2015a) in OpenSim (Delp et al., 2007). The model from Gritsenko et. al (Gritsenko et al., 2016) was modified to include separate digit segments representing an additional 16 degrees of freedom (DOFs). Here, we refer to the thumb as digit 1, index finder as digit 2, middle finger as digit 3, ring finger as digit 4, and little finger as digit 5. We did not include abduction (*abd*) / adduction (*add*) DOFs of the second to fifth digit metacarpophalangeal (MCP) and wrist joints. The resulting model simulated 18 DOFs in total (see Supplementary Tables). To improve the simulation of thumb movement, we added four thumb muscles, *opponens pollicis*, *flexor pollicis brevis*, *abductor pollicis brevis,* and *adductor pollicis*. Only a subset of muscles spanning the elbow, wrist, and hand joints were included in this validation, describing 33 musculotendon actuators representing 24 muscles of the distal arm and hand (Fig.1).

**Figure 1:**
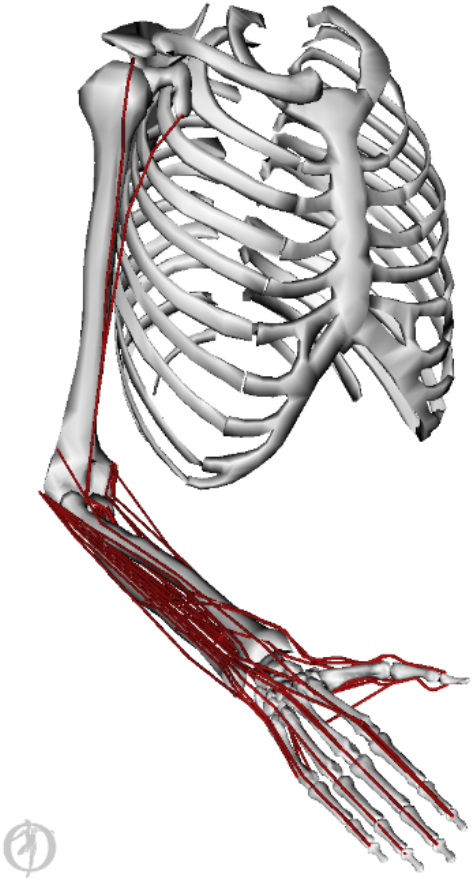
OpenSim upper-limb model of human muscles spanning distal arm and hand. The muscle paths (*red*) were defined relative to the skeletal landmarks and wrapping geometry.

Each musculotendon actuator in the model simulated active contractile and passive non-contractile forces generated by the muscles. The length-tension muscle property based on the Hill-type model simulated the active force generation (Zajac, 1989). The maximum active force for each actuator (*F*_*max*_) was set based on published physiological cross-section area (PCSA) (Chao et al., 1989) and muscle-specific tension (*σ*) = 35.4 (Arkin, 1941; Brand et al., 1981), so that *F*_*max*_ = *σ* · *PCSA*. All PCSA values originated from the same study (Chao et al., 1989) with the exception of *biceps*, which was not included in that study, so its parameters were derived from another study (Happee and Van der Helm, 1995). The maximum passive force component in the muscle model was empirically set to be 10% of the maximal force (*F*_*max*_). The simulated muscle properties were important for the functional validation of joint torques and the testing of constant moment arm assumptions described below.

### Datasets

Three datasets were created for the structural and functional validation of the MS model: *1*) published muscle moment arm measurements; *2*) published torque measurements in maximal voluntary contraction (MVC) tasks; *3*) simulated muscle moment arm values. The muscle moment arm measurements (Dataset 1) were used as a gold standard for the comparison with the simulated data (Dataset 3). The error between these datasets was reduced through the recurrent path adjustments described below (see Validation Process). Published torque data (Dataset 2) were used to validate ensemble force output from multiple agonist-antagonist muscles spanning the same joint. The goal here was to collect representative values, not to conduct a meta-analysis study.

### Dataset 1. Experimental Moment Arm Measurements

The experimental dataset of moment arm measurements was compared to the simulated relationships in Dataset 3 (below). We reviewed publications on moment arm measurements in human upper-limb and extracted their relationships with posture; however, these measurements were not of uniform quality. We selected sources with measurements in cadavers over other methods, e.g., simulated relationships, because these measurements were likely more reliable. If there were duplicate sources with similar methodologies, we selected the source with the most extensive information. Only seven sources were chosen to reduce the potential problems from combining multiple observations (Goislard De Monsabert et al., 2018). Then, the high-quality scans of published figures plotting muscle moment arms at the corresponding joint angle were digitized using specialized software (Rohatgi, 2018). The values of published moment arm and the corresponding DOF angle, termed muscle-DOF relationships, were expressed in SI units. Our search has not identified any moment arm measurements for muscles spanning the proximal and distal interphalangeal joints (PIP & DIP) of digits 3-5. The profiles of these muscle-DOF relationships were estimated using the published moment arms for the 2^nd^ digit. First, the ratios of moment arms at the 2^nd^ MCP joint and the 3^rd^ – 5^th^ MCP joints of homologous muscles were calculated. Second, these ratios were used to estimate the moment arms at the 3^rd^ – 5^th^ PIP and DIP joints from the 2^nd^ PIP and DIP moment arms of homologous muscles. For example, *extensor digitorum* moment arm profiles about the PIP joint of the middle finger (ED3) were copied from the index finger (ED2) and scaled by the ratio of moment arms for these muscles measured at the 2^nd^ and 3^rd^ MCP joints. Table 1 summarizes meta information for the included 81 muscle-DOF relationships (see Supplementary Materials for each muscle-DOF relationship and its source publication).

**Table 1:** List of sources with meta information that was included in the moment arm database.

| Source | Age (range), year | # Subjects | Measurement Method |
| --- | --- | --- | --- |
| (Haugstvedt et al., 2001) | 65 (41-90) | 8 (8M) | cadaver |
| (Bremer et al., 2006) | NP | 1 (1M) | cadaver and reconstructed molded model |
| (Loren et al., 1996) | NP | 5 | cadaver |
| (Koh et al., 2006) | 71 (50-90) | 11 (6M, 5F) | cadaver |
| (Fowler et al., 2001)* | 29 | 1 (1F) | MRI Scan |
| (Smutz et al., 1998) | 77 (72-84) | 7 (4M, 2F, 1 NP) | cadaver |
| (Gonzalez et al., 1997) | NP | NP | computer model (included for muscles without other data sources) |

\* Source indicated subject height 171.5 cm and weight 63.5 kg, M and F indicate male and female respectively, NP stands for not provided; MRI stands for magnetic resonance imaging.

### Dataset 2. Experimental Torque Measurements

Through a literature search, we found documentation of joint torques during maximum voluntary contraction tasks for 8 DOFs representing 6 hand joints. We used these experimental measurements to scale the maximum isometric force (*F*_*max*_) parameter in the Hill-type muscle model (Yakovenko et al., 2004; Zajac, 1989). This information was determined through the examination of a subset of all available studies from which four were selected based on the required experimental measurements at each DOF (Table 2). Critically, each isometric measurement was performed at a specific limb posture, which is expected to affect maximum force generation via muscle force-length relationship. We used an indirect estimate of MCP torque measurements based on the reported measurements of digit end-point forces and estimated moment arms (Shim et al., 2007). The estimated moment arms were different for each digit based on the placement of force sensor relative to the MCP joint in the experimental posture (7.5 cm digit 2, 8.0 cm digit 3, 7.5 cm digit 4, and 6.5 cm digit 5). These computed joint torques are marked with an asterisk in Table 2. Table 2 shows the summary of recorded values and the corresponding meta-information. These torque values are meant to be representative of an average person in their twenties.

**Table 2:** List of sources for torque measurements during maximum voluntary contraction tasks.

| Reference | Joint | Direction | Maximum Torque, Nm | Mean Height, m | Mean Age, year | Age Range, year | Weight, kg | Sex |
| --- | --- | --- | --- | --- | --- | --- | --- | --- |
| (Decostre et al., 2015) | Wrist | Flex<br>Ext | 14.5<br>11.4 | 1.77 | 25 | 20-29 | 74.9 | 27 M |
| (Gordon et al., 2004) |  | Pro<br>Sup | 4.64<br>4.38 | NP | 30.2 | 23.9-36.5 | NP | 11 M<br>3 F |
| (Bourbonnais and Duval, 1991) | 1<sup>st</sup><br>CMC | Flex / Add<br>Ext / Add<br>Flex / Abd<br>Ext / Abd | 4.2<br>3.4<br>2<br>1.6 | NP | 23.4 | 21-25 | NP | 12 F |
| (Shim et al., 2007) | 2<sup>nd</sup><br>MCP<br>3<sup>rd</sup><br>MCP<br>4<sup>th</sup><br>MCP<br>5<sup>th</sup><br>MCP | Flex<br>Ext<br>Flex<br>Ext<br>Flex<br>Ext<br>Flex<br>Ext | 3.15*<br>0.78*<br>3.36*<br>0.70*<br>2.1*<br>0.58*<br>1.59*<br>0.48* | NP | 22.5 | 20.5-24.5 | NP | 13 M<br>12 F |

\* estimated joint torque values; Flex = Flexion, Ext = Extension, Pro = Pronation, Sup = Supination, Add = Adduction, Abd = Abduction; NP = Not Provided, M = Male, F = Female.

### Dataset 3. Simulated Muscle Moment Measurements

The simulated muscle moment arm and length values were acquired from the OpenSim model in a uniform grid with 9 points spanning the maximal range of each DOF (Sobinov et al., 2019). This created 9^*d*^ unique postures per muscle, where *d* is the number of DOFs a muscle spans. The dataset contained muscle moment arms and length for all these postures. A muscle can span multiple DOFs (on average, 3), creating *d* moment arms per posture, *d*\*9^*d*^ total moment arm values per muscle. There was only one value of muscle length for a given posture, creating 9^*d*^ length values per muscle. Then, the total number of values for each muscle was 9^*d*^(2*d*+1), *e.g.*, an average muscle with *d*=3 was described by the sum of the following terms corresponding to 3*9^3^ postures with 3*9^3^ moment arms and 9^3^ muscle lengths for a total of 5,103 values. Figure 2 illustrates the dramatic increase in the number of postures for more complex muscles. The left panel shows a case for a single dimension with measured (grey) and some unmeasured (black) values within the physiological range. The middle panel is a typical representation of the two-dimensional case where a muscle spans two DOFs. The measurements are much sparser, rarely taken when both DOFs are manipulated. This problem of missing measurements worsens in the three-dimensional case, as in the right panel. Only a subset of postures in the three-dimensional domain contains measurements (no measurements are shown in this illustration). The most complex muscle is extensor pollicis longus (*EPL)* with *d*=6, which was described with 6,908,733 values.

**Figure 2:**
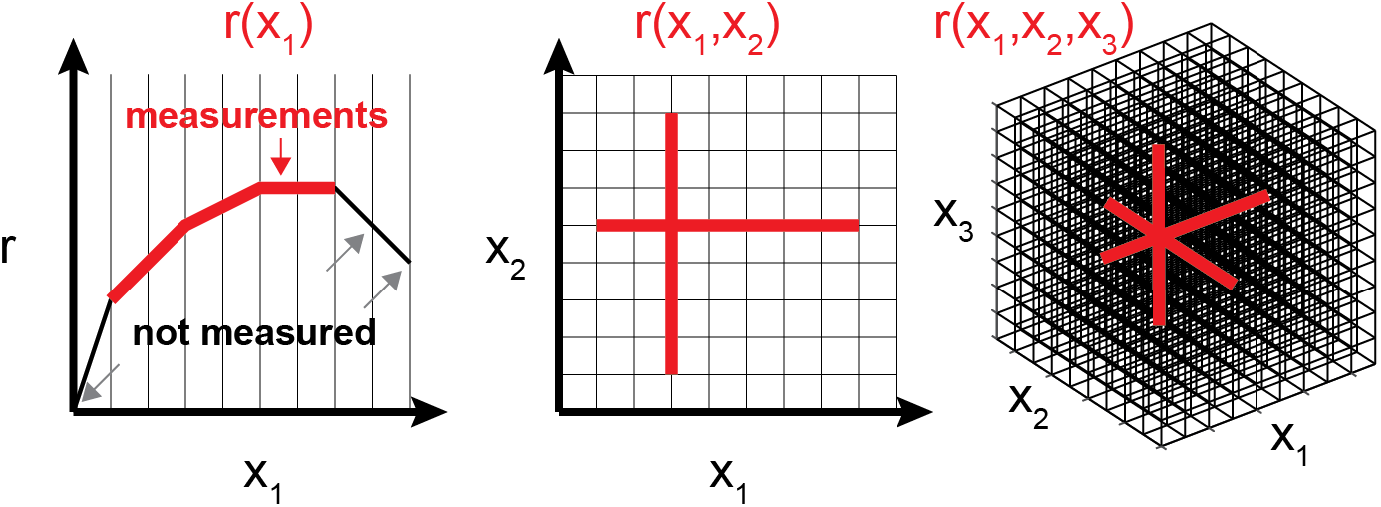
The problem of sparseness for musculoskeletal measurements. Several postures (~9) per DOF (x) are required to capture the profiles of muscle length and moment arm relationships (r). The number of values needed to be measured increase with additional dimensions actuated by each muscle, as illustrated by the increasing number of points from left to right panel. The vertices in the grid demonstrate the required measurements, and gray lines illustrate the typical data measured experimentally. The typical hand muscle spans over 3 DOFs requiring more than 729 independent measurements (right panel).

### Validation Process

We have identified three main types of errors in MS models: 1) errors in simulated muscle path for experimentally observed postures; 2) errors in simulated muscle path for experimentally unobserved postures; 3) force scaling mismatch between agonist-antagonist muscle groups. To mitigate these errors, we used a three-step process where the muscle paths were validated iteratively in the first two steps (see below), and then their ensemble behavior was validated in the third step.

### Step 1. Structural Validation for Experimentally Observed Postures

We evaluated the anatomical validity of musculotendon paths by comparing them to experimental measurements from Dataset 1. For each muscle, we initially set the origin and insertion points using a standard anatomical reference (Netter, 2011). The musculotendon paths were then repeatedly fine-tuned and checked to match the experimental and simulated moment arm values (Fig. 3a). This process involved manual adjustments of the wrapping objects and via-points of each muscle in OpenSim and comparing the changed moment arms to the Dataset 1. This dataset was updated every time a muscle path in the model was adjusted. For computational convenience, we used an accurate polynomial approximation of this dataset with low errors (<1%). The experimental and computed values were compared during structural validation and used in the estimation of forces (Step 3).

**Figure 3:**
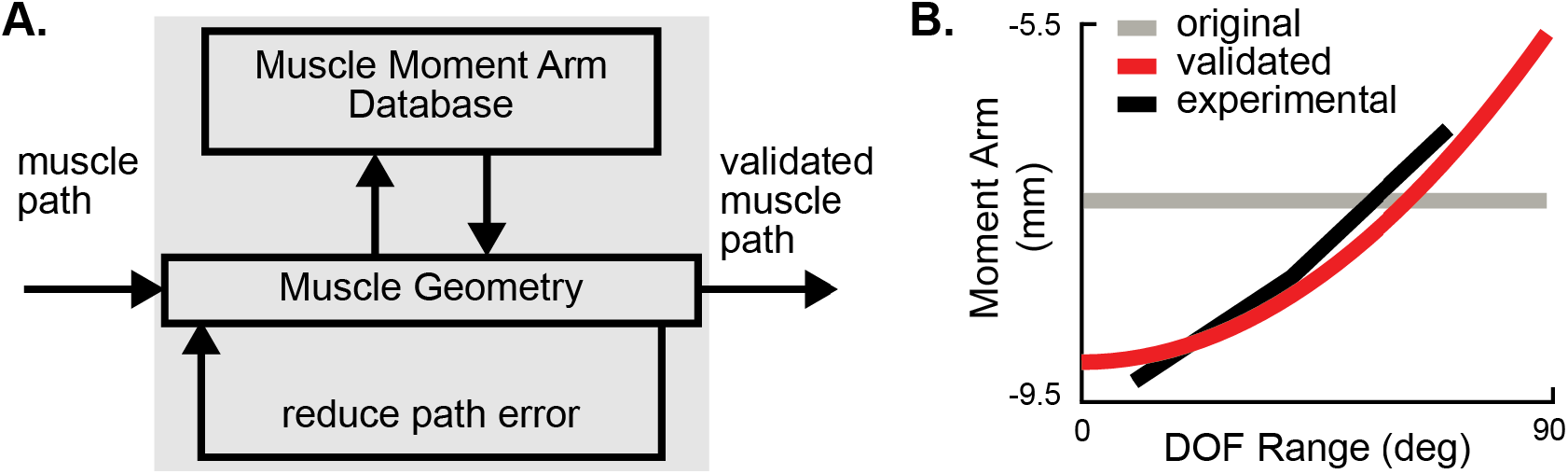
The process of structural validation using experimental data. A. Flow chart of moment arm validation. B. Example of simulated (before = gray, after = red) and experimental (black) moment arm profiles for the index finger *extensor digitorum* (ED2) muscle about the 2^nd^ MCP joint flex/ext DOF.

We used root mean square error (RMSE) and the correlation coefficient (*r*) between simulated and experimental values to determine the ‘good enough’ quality of moment arm profiles. ‘Good enough’ profiles had *r* > 0.7 and either RMSE < 1 mm or RMSE normalized to the range of experimental moment arm measurements < 0.4. The custom metric based on RMSE and *r* values expressed the operational definition of acceptable quality within our model. These values were chosen subjectively based on the examination of quality within the published model (Gritsenko et al., 2016; Saul et al., 2015a), and its performance in real-time decoding of motor intent (Boots et al., 2016; Mansouri et al., 2018). When these metrics were not met, indicating the low quality of simulated muscle paths, we interactively adjusted the muscle’s geometric constraints, via points and wrapping geometry, in OpenSim (Fig. 3a). The experimental data measured in human cadaver hands (Fowler et al., 2001) is closely represented by the simulated muscle-DOF relationship after validation (Fig. 3b). For example. to achieve this matched relationship for ED2 at the MCP joint, we moved incrementally a cylindrical wrapper off from the center of rotation for this DOF improving the path accuracy (Fig. 3b, gray to black). While this type of adjustment is straight-forward for simple DOFs, this process becomes time-consuming and challenging for muscles with interactions between several DOFs, e.g., *extensor pollicis longus* described by six-dimensional transformation. Many multidimensional muscle-DOF relationships have not been experimentally recorded and required additional error checks described in step 2.

### Step 2. Structural Validation for Experimentally Unobserved Postures

The goal of using biomechanical models for fundamental transformations of muscle activity into generated joint torques and valid limb motion requires data-driven model development. This task is challenging because experimental data for the description of moment arms in physiological postures is sparse. Moreover, the subjective examination of muscle characteristics in the high-dimensional space (*d*>3) of limb postures is limited. The following analyses were devised to resolve potential model failures, i.e., structural and functional discontinuities, in the posture space without experimental measurements:

*Zero-Crossing Error.* This analysis identified the functional discontinuity corresponding to a flip in the direction of muscle torque, which was indicated by the zero-crossing event in its moment arm profile. This type of failure could arise from the unaccounted interactions with wrapping objects in muscles spanning multiple DOFs. The testing for these crossing events was performed on each muscle-DOF relationship in Dataset 1. Zero-crossings occur when the global maximum and minimum have opposite signs for one or more DOFs that the muscle spans. Figure 4 shows an example with opposite signs of maximum and minimum values for *extensor carpi ulnaris* (ECU) in the original model. The superimposed profiles indicate changes in ECU wrist flexion-extension moment arms as a function of wrist flexion-extension angle in 9 wrist pronation-supination postures. The validated model had the extrema with the same sign after ECU was corrected from slipping off the geometric wrapper in the extreme wrist extension. All muscles were tested, and all zero-crossing events found were then examined in OpenSim for potential structural errors.
*Moment Arm Evaluation in Postural Extrema.* The goal of this analysis was to review the profile of moment arms for agonist-antagonist muscle groups at the postural extremes, i.e., postures corresponding to 0% and 100% of full physiological range of motion (ROM) for each DOF. The additional examination of moment arm dynamics in these postures was important because these locations often corresponded to failures in muscle wrapping geometry that, in addition to causing flipping in the sign of moment arms, could also lock the joint. This happened because of the imbalance between antagonistic muscles defined by the opposite signs of their moment arms in these postures. The imbalance was identified by the two-fold differences between the averages of peak-to-peak values for antagonistic muscle groups with moments around a particular DOF. These profiles were further scrutinized in OpenSim editor for the evidence of structural problems.

**Figure 4:**
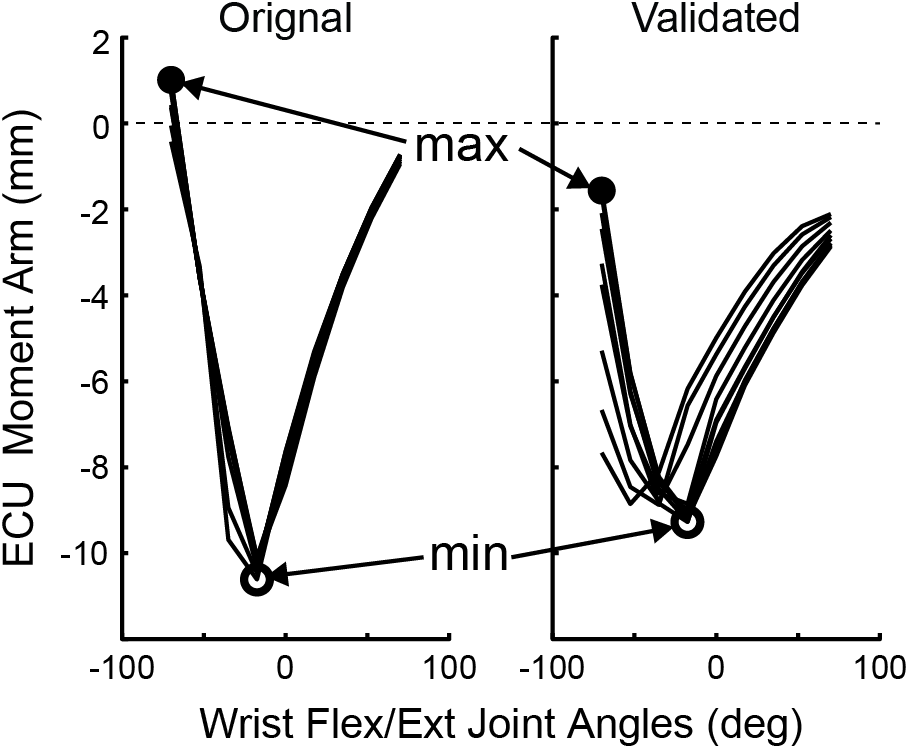
Example of structure validation for experimentally unobserved postures. ECU moment arm profiles for wrist flex/ext postures is shown for the original (left) and validated (right) models. The ECU muscle spans wrist flex/ext and wrist pro/sup DOFs, and its moment arm value depends on both joint angles, which is evident from the shown multiple profiles for varying pro/sup angle. The presence of an unexpected zero-crossing event when wrist was flexed from −50° indicated a potential error where muscle action flipped from extension to flexion moments in the original model. Zero-crossing events were examined and fixed through structural validation, as in the model shown in the right panel.

### Step 3. Functional Validation of Torque Generation

Data collation from multiple sources was expected to introduce incongruency between the measurements across multiple muscles, even though each muscle was consistent with a published observation (Goislard De Monsabert et al., 2018). To mitigate this problem across muscle groups, we validated joint torque generation relative to the torque in maximum voluntary contractions (MVC) at specific postures (see above, Dataset 2 Torque Measurements). We simulated the torque measurements in MVC trials by locking the model in the experimental posture and supplying the maximum activation to all muscles with the same sign of moment arms around the DOF of interest, i.e., agonists. The force generation was posture dependent and determined by a Hill-type muscle model (Hill, 1938; Yakovenko et al., 2004; Zajac, 1989).

The joint MVC torque was computed as the sum of all cross-products between muscle moment arms and forces (Fig. 5). The goal was to compute the scaling constant (*C*_*F*_) between experimental and simulated torques to adjust force generation across all muscles: 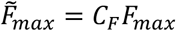. The adjusted 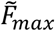 values were computed iteratively from distal to proximal DOFs. This approach locked the previous solutions for muscles with actions at distal DOFs and adjusted only the values of new (unlocked) muscles for each DOF. A single constant was assumed to scale parameters across all ‘unlocked’ agonists. For example, only *flexor digitorum, extensor digitorum, extensor indicis, and digiti minimi* muscles span MCP joints. Therefore, their 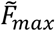 values were adjusted only based on toques about the phalangeal joints and remained the same (locked) during the validation of torques originating from more proximal muscles spanning wrist DOFs, *i.e.*, flexion/extension and pronation/supination. For the wrist DOFs, only the 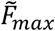 values of the muscles that do not span the phalangeal joints were adjusted to match wrist MVC, i.e. *extensor carpi ulnaris* (ECU), *flexor carpi radialis* (FCR), *flexor carpi ulnaris* (FCU), and *palmaris longus* (PL).

**Figure 5:**
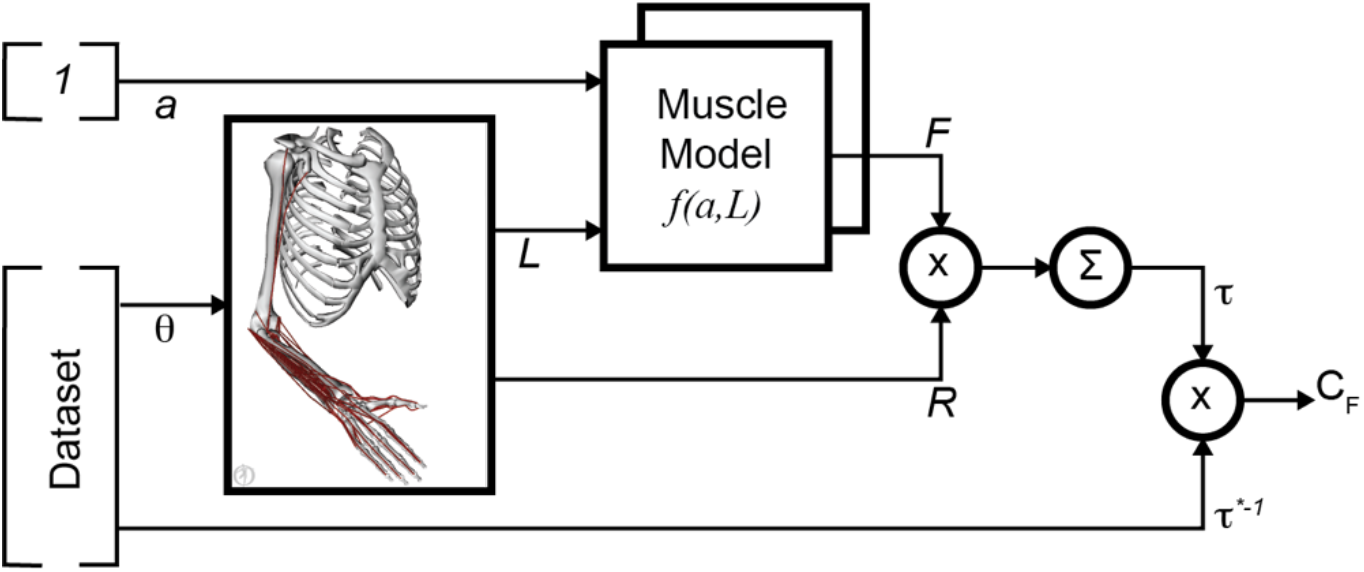
The process of scaling muscle force output. The torques at maximal voluntary activation (*a*) = 1 depend on joint posture (*θ*). *θ* were used to compute muscle length (*L*) and moment arms (R). Muscle force (*F*) was calculated from *a* and *L* using a muscle model for isometric conditions. The simulated torque (*τ*) was then calculated by summing the muscle force cross products with their moment arms (*R*) about each DOF. The scaling coefficient (*C*_*F*_) was computed using the simulated (*τ*) and the inverse of experimental torques (*τ*\*^−1^). The *θ* and *τ*\* values came from Dataset 2.

To validate the adjusted joint torques, we calculated the specific tension (*σ*) from the scaled model as 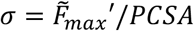 and compared it to experimental measurements and parameters in previous models. The experimental specific tension measures were obtained from human cadavers (Arkin, 1941; Brand et al., 1981; Haxton, 1944; Narici et al., 1988; Weijs and Hillen, 1985). The simulated specific tension values were obtained from (Buchanan, 1995) based on 4 musculoskeletal models (Amis et al., 1979; An et al., 1981; Edgerton et al., 1990; Murray et al., 1995).

### Statistical Analyses

All statistical measures used a significance value of α=0.05. The specific tension comparisons between simulated and experimental observations were performed by a one-way analysis of variance (ANOVA) with a Tukey post hoc test to determine which groups were different. The predictions of differences between constant and dynamic moment arms at postural extrema were tested with one-tailed t-tests with Bonferroni correction.

The comparison of dynamic relationships between moment arms was performed using hierarchal clustering of correlation matrix (Gritsenko et al., 2016). First, a correlation matrix was computed between the moment arm relationships across all muscles per DOF. Then it was transformed into the heterogeneous variance explained (HVE) metric defined as *1-r*^*2*^ for *r*>*0* and *1*+*r*^*2*^ for *r*<*0*. This metric assigned short distances close to 0 to positively correlated moment arm profiles and distances close to 2 to negatively correlated moment arm profiles, weak correlations were assigned an intermediate distance close to 1. We then used hierarchal clustering of HVE to define muscle groups based on their moment arm profiles. We used this grouping to further evaluate the magnitude of errors that may be caused by the assumption of constant moment arm profiles.

With the development of ever more challenging complex musculoskeletal models the use of constant moment arms across all postures is a convenient approach that partially mitigates structural errors. However, the trade-off related to the error in the estimation of torque due to the coordination of musculoskeletal relationships remains unclear. We will test the following specific predictions about the coordination of wrist muscles to evaluate the importance of non-linear changes in the moment arm profiles:

1. The dynamic moment arms are smaller than the constant moment arms when the muscle length is long.
2. The dynamic moment arms are larger than the constant moment arms when the muscle length is short.

## Results

In this study, we developed a validation process for the structure and function of a detailed arm and hand musculoskeletal model. We used a combination of quantitative and qualitative metrics to correct anatomical errors during the process of adjusting muscle paths relative to the skeletal landmarks to fit available experimental relationships. Then, we applied our quantitative analysis to identify two possible muscle path errors in the absence of experimental observations: *i*) the zero-crossing error due to the flipping of moment arm sign changing muscle function, *ii*) inadequate force generation in extreme postures. Each instance of tentative errors identified with these methods was examined and manually corrected, if necessary. These structural corrections were followed by the adjustment of balance in force generation between agonistic muscle groups using experimental MVC measurements. We used the adjusted model to compare experimental and simulated muscle-specific tension values and to evaluate the potential discrepancies in torque calculations with the assumptions of constant or variable muscle moment arms at multiple wrist postures.

### Validation Process

The structural quality of the MS arm model increased after muscle path validation. We assessed five types of errors in muscle paths for 33-musculotendon actuators controlling 18 DOFs of the arm and hand (Fig. 1). Each muscle spans on average 3 to 4 DOFs with EPL spanning the maximum of 6 DOFs in this model, as shown in columns of original (left) and validated (right) panels with categorical representations. The five categories describe dynamic and static profile errors ranging from the worst (red) to the best (green, Fig. 1B). The ‘good’ category (green) was associated with the least amount of error between the experimental (Dataset 1) and simulated (Dataset 3) moment arm measurements.

The highest quality (green) category was assigned to muscles with the high correlation between experimental and simulated profiles of moment arms (*r*>0.72), with an average of *r*=0.92±0.07 (s.d.) after validation, and with the minimal static offset that was less than 5% of the range from minimum to maximum. The yellow category included profiles with the high offset error. The *pink*, *orange*, and *yellow* categories represent errors that do not include the zero-crossing error, which identifies the *red* category. Errors in the original model from Gritsenko et al. (Gritsenko et al., 2016; Saul et al., 2015a) had multiple failures represented by the flipping sign of moment arms (*red* category). Moreover, the number of muscles in the top category increased from only 8.6% in the original model to 73.5% in the validated model, and the majority of profiles showing the incorrect zero-crossing locations (*red* category) were corrected (Fig. 6C). Although not all moment arm profiles were assigned to the top category, all of them were improved through the validation process. The overall improvement of model quality was quantified with a metric that assigns progressive scores to each color (*red*=1, *pink*=2, *orange*=3, *yellow*=4, and *green*=5). The total score was calculated only for muscles present in both models. The original model score was 277, and it was 418 for the validated model, which indicates 1.51 times improvement on this ordinal scale.

**Figure 6:**
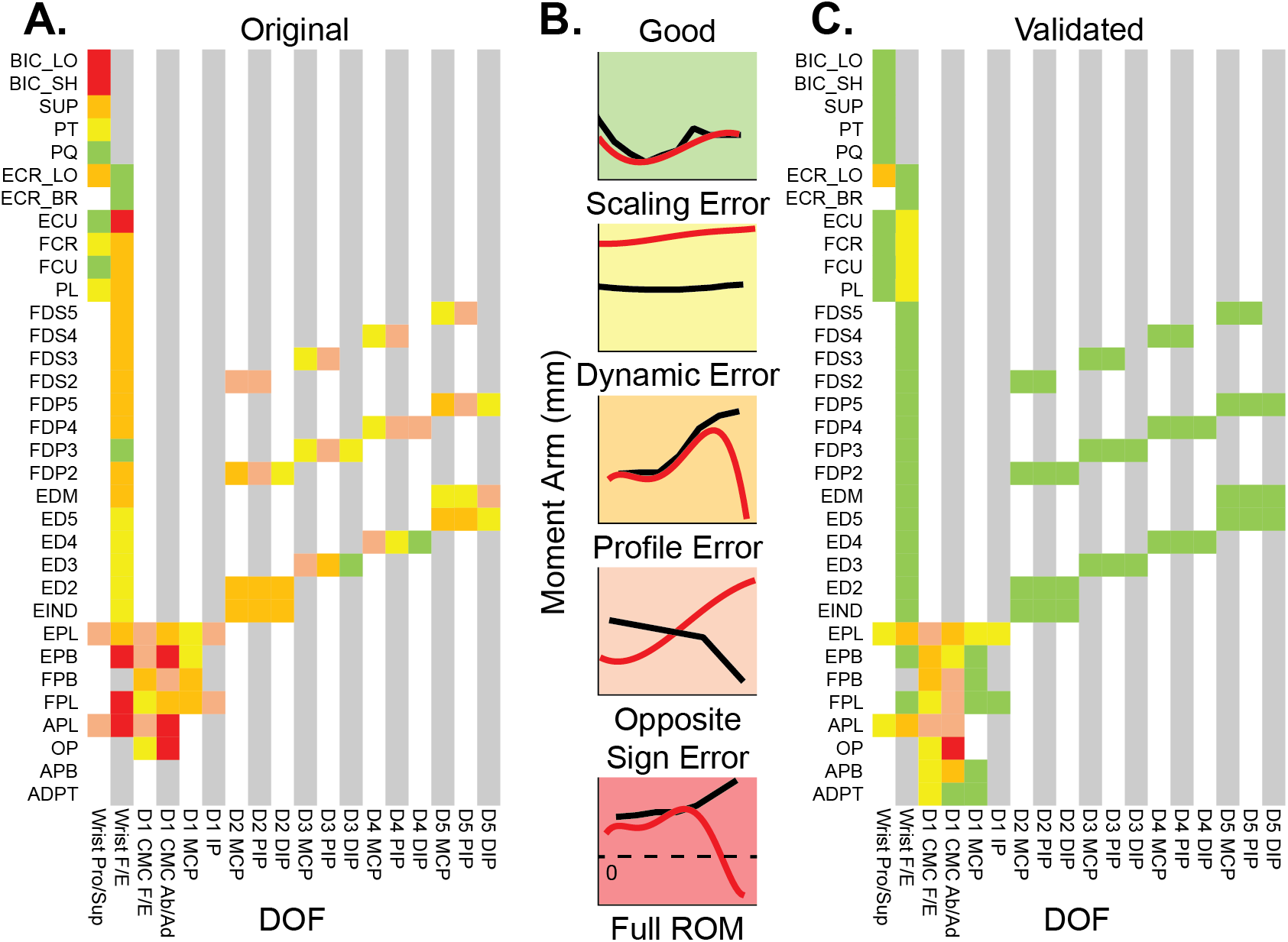
The comparison of structural quality between models with and without structural validation. A. Categorical errors in the original model from Gritsenko et al. (Gritsenko et al., 2016; Saul et al., 2015a) for all DOFs (column labels) and all muscles (row labels). Missing values in the grid represent DOFs that a given muscle does not span. F/E indicated flexion/extension; Pro/Sup indicates pronation/supination; Ab/Ad indicates abduction/adduction. B. Examples of errors in each category. C. Categorical errors in the validated model after the manual path validation.

The validation procedure did not fully correct some profile errors in 8 thumb and 5 wrist muscles. These high dimensional relationships were difficult to correct manually using the iterative process described in Step 1 and may require an automated optimization in the future work.

The experimental dataset sparsely represented all the muscle-DOF relationships. The unobserved domain was examined for possible zero-crossing errors. We found that the original model had multiple zero-crossing instances. The zero-crossing instances were distributed uniformly within ROM (Fig. 7A). Each instance of zero-crossing in the original model was examined and iteratively corrected only if the crossing occurred due to problems with wrapping geometry. The flip of function in muscles spanning wrist pronation/supination and CMC flexion/extension DOFs was supported by the published experimental profiles included into our dataset (Fig. 7B).

**Figure 7:**
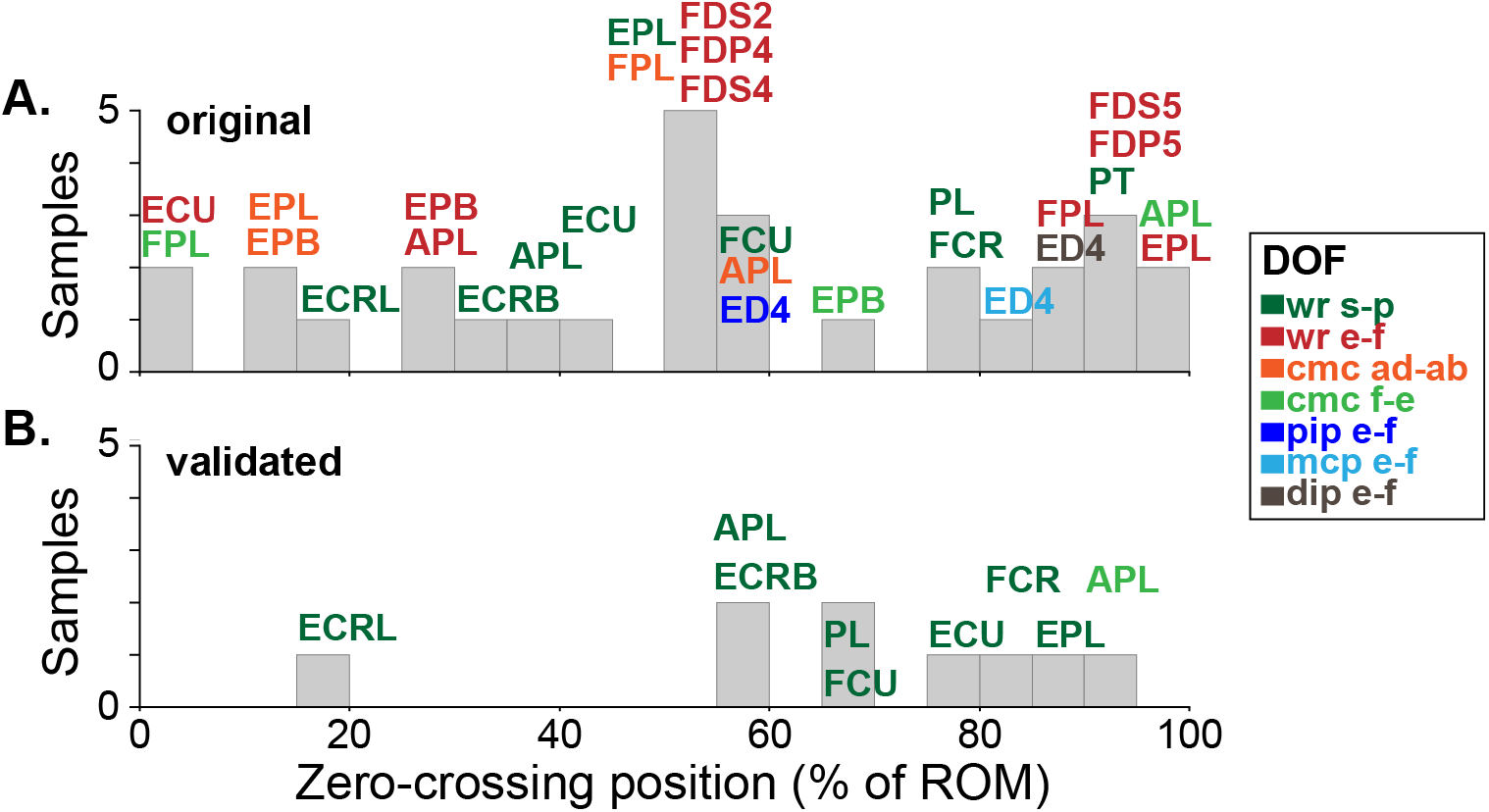
Histograms of muscle moment arm zero-crossing events as a function of position. The colors represent the DOFs (described in the legend) where the crossing takes place. s-p indicates supination/pronation; e-f indicates extension/ flexion; ab/ad indicates abduction/adduction. The distributions are shown for original (A) and validated (B) models.

Next, we validated the model force generation using the MVC values (Dataset 2, **Table 2**) for digit and wrist DOFs. The values for the *F*_*max*_ parameter in the Hill-type muscle model were adjusted in isometric simulations for specific postures corresponding to Dataset 2, as described in Step 3. We used the *distal-to-proximal* parameter adjustment method to minimize errors across multiple joints. The original estimates of *F*_*max*_ values from the published models produced high errors in the wrist muscle torques (Fig. 3A, red circles). The functional validation scaled down the *F*_*max*_ of finger muscles, which then, in turn, led to the adjustment of the *F*_*max*_ for wrist muscles to account for the force being produced by the finger muscles at the wrist. This process generated a physiologically relevant maximum torque at the wrist and 2^nd^ – 5^th^ digits (Fig. 8A, blue circles). The moment arms of complex thumb muscles were not fully corrected (shown in Fig. 1C) and that limited the validation of torques for thumb joints.

**Figure 8:**
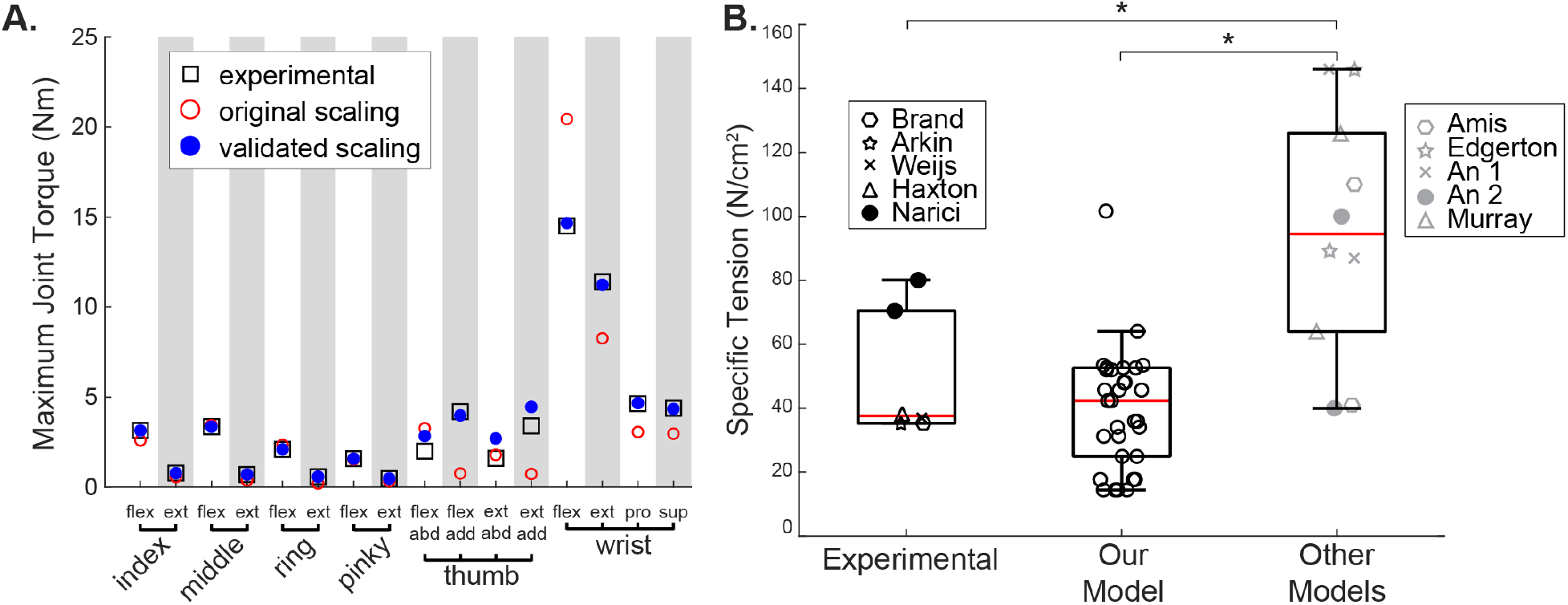
Functional validation of joint torques and muscle forces. A. The torque profiles were reconstructed for eight arm and hand DOFs. The *black* squares are the published maximum torque values, the *red* circles are the torque values simulated with a Hill-type muscle model using the original muscle paths, and the *blue* circles are the torque values after muscle path validation. B. Muscle-specific tension values. The published experimental data (Experimental), data simulated with the validated model (Our Model), and other simulated published data (Other Models listed in the legend) were compared. The asterisk marks significant differences identified by post-hoc tests.

**Figure 9:**
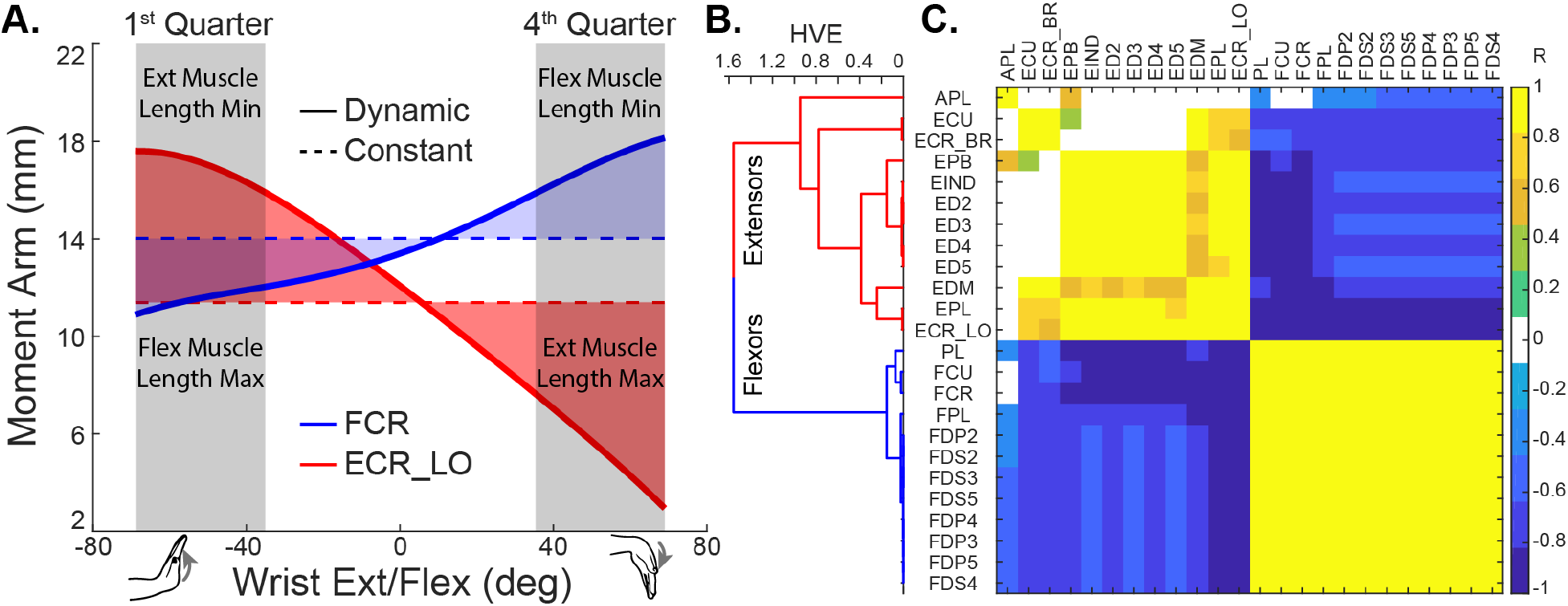
The comparison of constant and dynamics moment arms. A. Example wrist flexor (blue) and extensor (red) moment arm profiles around wrist flexion/extension DOF. B. The dendrogram produced by the hierarchal cluster analysis of HVE obtained from the correlation matrix in (C). C. The correlation matrix of *r*-values between pairs of muscle moment arm profiles for a wrist flexion/extension DOF.

To test the validity of scaling the maximum force values to the experimentally observed maximal joint torque values, we calculated the muscle specific tension values that emerge after the functional validation. ANOVA showed that the experimental and simulated muscle specific tension values are different (Main Effect of Group: f=6.54, p=0.032). Tukey post-hoc tests showed that the adjusted muscle specific tension values in our validated model were not significantly different from the experimentally measured values, while the simulated values from the “other models” were significantly higher than both the experimental values and our simulated values (Fig. 3B).

### Constant vs. Dynamical Moment Arms

We evaluated the error that could arise from the use of constant moment arm assumption for muscles actuating the wrist. In general, The moment arm profiles potentiate wrist torque generation. For example, *flexor carpi radialis* (FCR) wrist flexor moment arm increases with the increase in wrist flexion (Fig. 4A). Similarly, the moment arm of wrist *extensor carpi radialis longus* (ECRL) increases with the increase in wrist extension. As muscles shorten their ability to generate force decreases, but the increase in their moment arms maintains similar capacity for torque generation. These general relationships were homologous across wrist flexors and extensors, as shown by the correlation analysis of moment arms in Fig. 4B&C. The hierarchal clustering analysis showed close similarities in muscle-DOF relationships among flexors and among extensors while separating the flexors from extensors (mean HVE≈1.6, Fig. 4B). Next, we compared the differences of moment arm averages in extreme postural quartiles (dynamic condition, the solid line in Fig. 4A) against the averages across the full ROM (constant condition, dashed line in Fig. 4A). We found that the dynamic moment arms are significantly smaller than the static moment arms by mean 15.0 ± 14.8% (s.d. across DOFs) when the muscles are long and *vice versa* by 19.9 ± 14.2% when the muscles are short (p<0.0001 for both tests). Thus, substituting dynamic moment arm profiles with constants would result in a mean decrease of 34.9% in the amplitude of wrist torque generated for a full range of motion movement.

## Discussion

In this study, we developed a novel model validation method that improves the anatomical and functional accuracy of MS models. The process improves the structural path accuracy across the full range of all 18 DOFs using the multidimensional database of experimental measurements, skeletal landmarks, and the detection of potential structural errors. The functional validation process then relies on the corrected muscle moment arms to scale muscle forces to match the maximum torques recorded in the literature (see Datasets). Scaling the forces was vital because it adjusted the torque values of the model to stay within the biological boundaries of torque output. As a result of this validation process, we have improved the accuracy of the previous musculoskeletal model and tested the importance of nonlinearities in moment arm profiles.

We found that the assumption of constant moment arms for wrist and digit muscles can lead to about 35% error in the estimation of wrist flexion-extension joint torque. The moment arms near the extremes (Fig.7A, grey shaded regions) varied, on average, approximately 17±14% from the constant moment arm values. These values are supported by the previous demonstration of force generation in different wrist postures (Gonzalez et al., 1997). This means that the model with constant moment arms will grossly underestimate the muscle forces produced at postural extrema for a given level of muscle recruitment (Crouch and Huang, 2016); however, model simplifications and the use of lumped parameter assumptions may outperform other applications using inaccurate models. We provide further computational evidence for the importance of musculoskeletal organization in the estimation of forces with the use of improved models with the high-dimensional validation described in this study. The dynamic moment arm profiles were crucial for reproducing the measured maximal isometric torque in different postures of the human hand. Similarly, we expect that these structural details will be important in the decoding of motor intent from EMG in prosthetic or orthotic applications where MS properties contribute to the estimation of state-space variables for the control system (Menegaldo, 2017).

Musculoskeletal models are growing in complexity from low-dimensional, yet insightful, models (Blickhan, 1988; Full and Koditschek, 1999) to much more complex and detailed (Seth et al., 2018). Accurate implementation is ever more challenging partly because the appropriate data is extremely sparse. Consequently, the validation of model parameters across the physiological domain of postures or explicit limitations are difficult to evaluate and describe (Thacker et al., 2004). The typical validation domain spans measurements in a limited range of postures, usually not exceeding single-DOF variations and not often covering the full ROM of the joints a muscle actuates. Fig. 2B illustrates conceptually the sparseness of experimental data for a single muscle relative to the domain of interest. This constitutes a major limitation for the current models, including this work. To improve the accuracy of our model, we collected a large dataset of muscle-DOF relationships. However, the published data had two major limitations. The first limitation was that moment arms were measured using different methodologies for subsets of several muscles from people of different ages and genders causing discrepancies in the measurements. The second limitation was that the moment arms were, by necessity, measured for a subset of physiological postures and DOFs. To overcome these limitations, we developed a validation process that evaluated the moment arm values in all possible postures and found anomalies that could jeopardize model performance. These anomalies tended to occur at the extremes of ROM, because these postures were associated with sharp angles between bones, where wrapping objects were often unable to constrain appropriately the muscle paths (example in Fig.4). All the structural anomalies in muscle-DOF relationships ranging from inaccurate offset to sign flipping were identified by our validation process and corrected, with the exception of the opponens pollicis (OP) moment arm about the CMC abd/add DOF (Fig. 6). The validation of the moment arms about the thumb posed the biggest challenge due to the complexity of movement of the two-DOF CMC joint and the anatomical complexity of all thumb muscles spanning multiple joints. The validation process successfully identified the limitations of moment arm simulation about the CMC joint, so that is will be possible to overcome them in the future.

The functional validation process was crucial for enabling the physiologically relevant performance of the model by ensuring that the joint torques do not exceed physiological values. This process overcame two main limitations of sparse published data: 1) inconsistencies in the measured moment arm dataset and 2) inaccuracies due to the simplified simulation of muscle properties. In particular, inaccurate estimates of the maximum force (*F*_*max*_) a given muscle could lead to unphysiological solutions for muscle-specific tension. The muscle-specific tension in the validated model was in general agreement with published experimental values and significantly improved relative to in previous models (Fig.8B). These results further support our conclusions that the structural validation followed by functional validation improves the physiological accuracy of MS models.

As an outcome of the validation process, we have developed a generic arm model that is suitable for use in biomimetic control applications and for the biomechanical analysis of multiple types of arm movement. The model was validated to represent an average young adult by having its torque output and muscle paths match published average data. The model can be scaled to individual body size by scaling the size and mass of each segment to anatomical proportions based on individual height and weight (Winter, 2009). However, functional validation will need to be repeated to account for any changes in the muscle model parameters, such as *F*_*max*_ values, pennation angles, or *PCSA* values. The moment arm and muscle length values from the validated model can also be extracted for real-time applications (Sobinov et al., 2019), which provides a valuable resource for human-machine interfaces.

## Conclusions

In conclusion, we developed a validation process that can be used to create robust and physiologically accurate MS models. We applied this process and validated an MS model of the human forearm. We have also evaluated the impact on torque generation for the common computational assumption of constant moment arms for muscles actuating wrist joint. These results came from our validation of both the anatomical structure of the model and the functional torque output. Therefore, both structural and functional validation procedures are important for the development of generalizable models.

## Supplementary Material

The supplementary materials include PDFs of all the muscle-DOF profiles from experimental and simulated datasets that were used to create Figure 6. It also includes an Excel file for each muscle-DOF describing the experimental data profile and its source.

